# A framework for large-scale dynamic metabolome drug profiling in mammalian cells: a case study analysis of the anti-cancer drug dichloroacetate

**DOI:** 10.1101/250670

**Authors:** Sébastien Dubuis, Karin Ortmayr, Mattia Zampieri

## Abstract

Metabolic profiling of cell line collections have become an invaluable tool to study disease etiology, drug modes of action and personalized medicine. However, large-scale *in vitro* dynamic metabolic profiling is limited by time-consuming sampling and complex measurement procedures. By adapting an MS-based metabolomics workflow for high-throughput profiling of diverse adherent mammalian cells, we establish a technique for the rapid measurement and analysis of drug-induced dynamic changes in intracellular metabolites. This methodology is scalable to large compound libraries and is here applied to study the mechanism underlying the toxic effect of dichloroacetate in ovarian cancer cell lines. System-level analysis of the metabolic responses revealed a key and unexpected role of CoA imbalance in dichloroacetate toxicity. The herein proposed strategy for large-scale drug metabolic profiling is complementary to other molecular profiling techniques, opening new scientific and drug-discovery opportunities.

## Introduction

A major bottleneck in drug discovery pipelines is the lack of mechanistic information on the targets of selected lead compounds. Large-scale approaches enabling the characterization of cell responses to external perturbations have therefore turned into highly relevant technologies in drug discovery and development^1–4^. Among these approaches, the systemic profiling of drug-induced changes in model organisms at the mRNA and protein level^5,6^ has provided invaluable insights into drug modes of action (MoA)^7–9^, drug-drug interaction mechanisms^10^ and drug repurposing^2,11^. Conceptually similar to transcriptomics and proteomics platforms, metabolomics provides an orthogonal multi-parametric readout aiming at quantifying the full spectrum of small molecules in the cell, the so-called metabolome. Applied to drug discovery research, metabolome profiling of drug-perturbed cell lines *in vitro* was key in revealing drug MoA and in identifying potential weaknesses in cellular drug response as well as genetic polymorphisms associated with drug susceptibility^12–21^.

Metabolomics-based approaches have a notable advantage over existing functional genomics platforms in that they enable an unparalleled throughput^22,23^. However, despite significant advancements in high-resolution mass-spectrometry profiling of cellular samples^23–26^, efficient experimental and computational workflows for large-scale dynamic metabolome profiling in mammalian cells *in vitro* are lagging behind. Metabolome screenings that adopt classical metabolomics techniques^27,28^ are often hampered by a limited throughput, laborious sample preparation and the lack of rigorous, yet simple, data analysis pipelines to interpret dynamic metabolome profiles. To address these limitations, our group developed a high-throughput and robust method to perform large-scale static metabolic profiling in adherent mammalian cells^29^, using a 96-well plate cultivation format combined with time-lapse microscopy and flow-injection time-of-flight mass spectrometry^25^ (TOFMS). Here, we extend this methodology to allow rapid sample collection and the analysis of dynamic changes in the intracellular metabolome of diverse mammalian cell lines upon external perturbations. We applied this methodology to profile the diversity of metabolic adaptive responses in five ovarian cancer cell lines to an activator of pyruvate dehydrogenase (i.e. dichloroacetate) and a lactate dehydrogenase inhibitor (i.e. oxamate).

The presented framework for *in vitro* large-scale dynamic metabolomics of perturbed adherent mammalian cell lines is complementary to and scales with high-throughput growth-based phenotypic screens of large compound libraries. Moreover, we provide a proof of principle that our approach can characterize compound MoAs, and can generate testable predictions to elucidate the origin of drug response variability. Such a platform may complement and improve the translational value of classical *in vitro* phenotype-based drug screenings^30^, and provide insights on the mechanisms of action of small molecules facilitating early stages of drug discovery^31–33^.

## Results

### High-throughput dynamic metabolic profiling of drug action in mammalian cell lines

Large-scale metabolic profiling of transient drug responses among diverse cell types necessitates new methodologies enabling parallelized and rapid sample collection, high-throughput metabolic profiling and an effective normalization approach for metabolomics data. Here, we developed a combined experimental-computational approach enabling the rapid profiling of dynamic changes in the baseline metabolic profile of diverse cell lines in parallel. This approach was applied here to study the metabolic responses of five ovarian cancer cell lines to two enzyme inhibitors: dichloroacetate and oxamate.

In brief, the five different cell lines IGROV1, OVCAR3, OVCAR4, OVCAR8 and SKOV3 were grown in parallel in 96-well plates for four days. Cells were exposed to the corresponding drug dose yielding 50% growth inhibition (*GI_50_*) (Table 1) and metabolomics samples were collected every 24 hours, following the extraction protocol described in ^29^ and summarized in Supplementary Figure S1. In the present study, nine replicate plates were prepared: one plate served to continuously monitor cell growth via cell confluence by time-lapse microscopy using an automated multi-well plate reader (Figure 1), while the remaining plates are used for metabolome extraction immediately before, and at 24, 48, 72 and 96 hours after drug exposure (Supplementary Figure S1). At each sampling time point, one plate was used to generate cell extract samples, while the second plate served to determine extracted cell numbers per well using bright-field microscopy^29^. Cell extract samples were profiled by flow injection analysis and time-of-flight mass spectrometry (TOFMS) as described previously^25^, enabling high-throughput analysis of large sample collections. The detected ions were annotated based purely on the accurate mass, and by assuming that deprotonation is the most frequent and reliable form of ionization in negative mode. By matching measured m/z against calculated monoisotopic masses of metabolites listed in the Human Metabolome Database (HMDB^34^) and in the genome-scale reconstruction of human metabolism (Recon 2^35^), we putatively annotated 2482 ions (Supplementary Data Table S1). Importantly, in absence of prior chromatographic separation, FIA-TOFMS cannot distinguish isobaric metabolites as well as in-source fragments that are detected at the identical exact mass.

**Figure 1.**
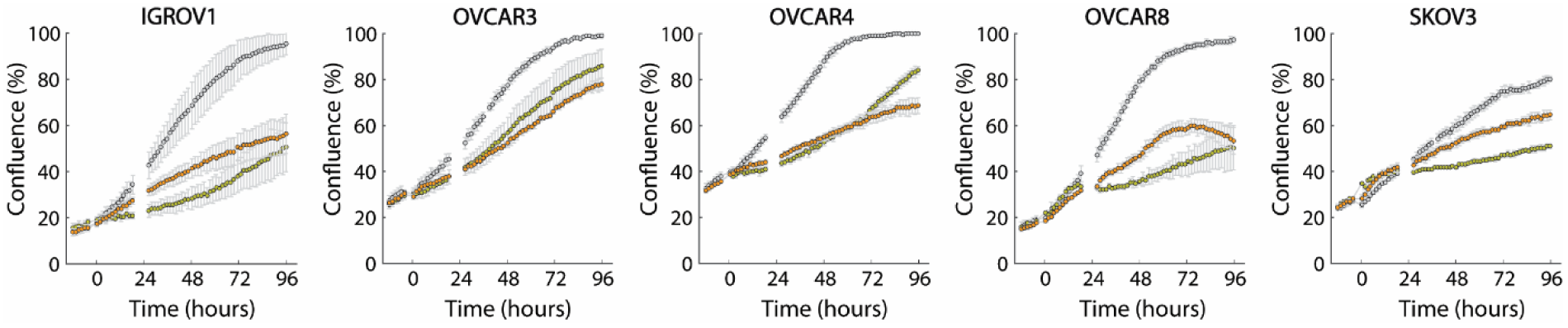
Growth of ovarian cancer cell lines upon drug perturbation. Cell confluence measured by time-lapse microscopy during growth of five ovarian cancer cell lines in RPMI1640 medium (i.e. untreated condition in grey), and upon dichloroacetate (orange) and oxamate (green) treatments (Supplementary Figures S10). Cell confluence is reported as mean ± SD across three replicates.

**Table 1.** Drug concentrations used in metabolomics experiments. The given concentrations correspond to the effective concentration in the medium.

| Cell line | Oxamate dose | Dichloroacetate dose |
| --- | --- | --- |
| IGROV1 | 13 mM | 11 mM |
| OVCAR3 | 40 mM | 12 mM |
| OVCAR4 | 18 mM | 25 mM |
| OVCAR8 | 6.7 mM | 25 mM |
| SKOV3 | 31 mM | 25 mM |

To estimate time-dependent (e.g. drug-induced) changes of intracellular metabolites from non-targeted metabolomics data we employed a regression-based analysis strategy. This approach compares transient changes in the metabolome of drug-treated cells against steady-state unperturbed cell metabolic profiles (Supplementary Figure S2). Steady-state metabolic profiles of unperturbed cells were determined following the approach described in ^29^, which is here briefly summarized. By definition, the intracellular concentrations of metabolites at steady state are constant in time. Hence, in samples from unperturbed growing cells for each metabolite *i* in cell line *j*, measured intensities, I_j,i_, scale proportionally with the metabolite abundance in the cell [m_i_], times the extracted cell number (N_c_, derived from bright-field microscopy^29^, see Supplementary Figure S3):

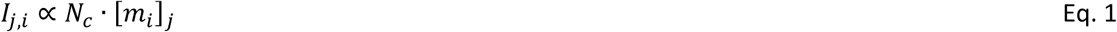

Hence, we can model the measured metabolite intensities in a given cell line as follows:

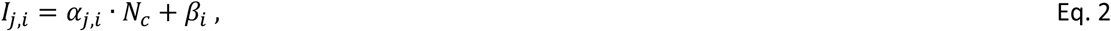

where β_i_ is an offset value representing the experimental MS background signal, and α_j,i_ represents the abundance of metabolite *i* per cell. Notably, α_j,i_ contains an unknown scaling factor that is reflective of the fundamental proportionality between metabolite concentration and MS signal intensity. For each metabolite, we use a multiple regression scheme to fit the linear model and regress the cell line-specific α values and the offset β across all cell lines at once (Supplementary Figure S4). To assess the reliability of the parameter estimates, each fitted parameter is associated with a p-value (F statistic with the hypothesis that the coefficient is equal to zero). It is worth noting that this procedure allows systematically filtering out: (i) annotated ions which are unlikely to originate from extracted cells, because the measured ion intensity does not exhibit any dependency with the cell number, and (ii) ions for which the measured intensities are below the detection limit, and the estimated cell line-specific α values are below or close to 0. Out of the 2482 ions annotated in the ovarian cancer cell dataset, we obtained relevant parameter estimates for 1546 putatively annotated ions, i.e. α > 0, and p ≤ 0.001 in at least one cell line, with a median coefficient of variation of 17.4% (Supplementary Figure S5).

To evaluate dynamic metabolite changes upon an external perturbation, we calculate time-dependent fold-change values for each metabolite based on the parametrized model derived from steady-state unperturbed metabolome profiles. For each metabolite *i* and time point *t* after exposure to treatment *D,* we estimate the deviation of metabolite concentration from the unperturbed steady-state condition as follows:

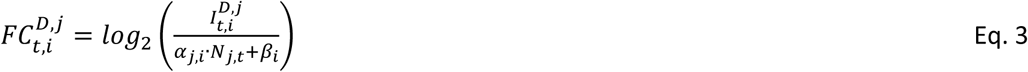

where 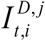 is the intensity measured for metabolite *i* at time *t* after exposure of cell line *j* to compound *D*. 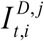 is compared to the corresponding theoretical steady-state unperturbed metabolite intensity (denominator in Eq. 3) which is calculated from the previously estimated *α* and β parameters and the extracted cell number at the time of sampling, *N_J,t_,* derived from bright-field microscopy images (Supplementary Figure S1 and S3). For each time point, the difference between measured and expected metabolite intensities is expressed in log*_2_* fold-changes, and significance is quantified by means of p-values from t-test analysis. In the following, our metabolome profiling pipeline was applied to investigate the metabolic response to two small-molecule agents, dichloroacetate and oxamate (Figure 2a-b), in the five ovarian cancer cell lines: IGROV1, OVCAR3, OVCAR4, OVCAR8 and SKOV3.

**Figure 2.**
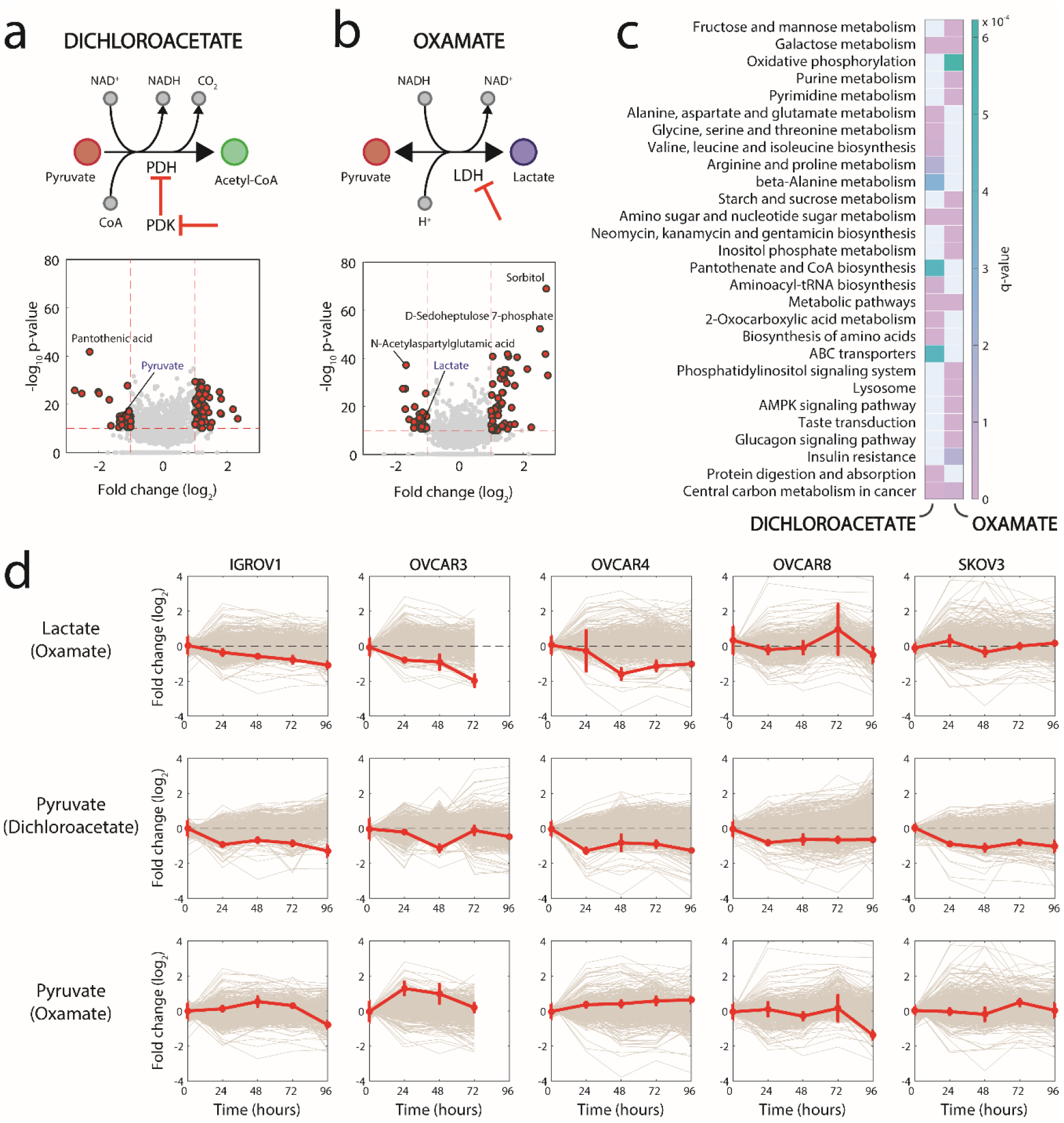
Analysis of transient dynamic metabolic changes upon external perturbation. **(a-b)** Schematic representation of the enzymatic reactions targeted by oxamate and dichloroacetate, and volcano plots summarizing overall metabolome changes. Each dot corresponds to a metabolite. The – log_10_ of the product between minimum p-values over the time course across the five cell lines is plotted against the median of maximum fold-changes. Metabolites highlighted in red have an absolute log_2_ fold-change > 1 and a p-value ≤ 1e-10. **(c)** KEGG pathway enrichment of metabolites consistently affected by dichloroacetate and oxamate treatments, highlighted in panel **b**. Only pathways with a q-value ≤ 0.001 are considered. **(d)** Time-dependent fold changes (red line) of lactate upon oxamate treatment, and pyruvate upon oxamate and dichloroacetate treatments. Data are the mean ± S.D. of three replicates. The profiles of all other detected metabolites are shown in grey.

### A role for coenzyme A metabolism in mediating the cellular response to dichloroacetate

Dichloroacetate (DCA) is a mitochondria-targeting small molecule that activates pyruvate dehydrogenase (PDH) by inhibiting pyruvate dehydrogenase kinase (PDK)^36^. By blocking PDH phosphorylation in mitochondria, dichloroacetate favors an increased production of acetyl-CoA from pyruvate and coenzyme A (CoA). To date, the exact mechanism by which dichloroacetate is toxic to cancer cells has remained unclear^37^. Recent studies suggested that the activation of PDH diverts metabolism from fermentative glycolysis to oxidative phosphorylation, leading to a loss in mitochondrial membrane potential and a reopening of voltage‐ and redox-sensitive mitochondrial transition pores, which ultimately triggers an apoptotic cascade in cancer cells^36^.

Despite large differences in doubling times (Supplementary Figure S6), degrees of invasiveness^38^ and metabolic phenotypes^39^ (Supplementary Figure S7), the five selected ovarian cancer lines exhibit common metabolic adaptive changes to dichloroacetate exposure. In our dynamic metabolome data, we observed (i) a consistent reduction in pyruvate levels across all cell lines upon dichloroacetate treatment (Figure 2d), (ii) a reduction in lactate secretion (Supplementary Figure S8), and (iii) a significant increase in the total pools of CoA and acetyl-CoA (Figure 3a and Supplementary Figure S9). Strikingly, the largest metabolic change across all five cell lines was a marked depletion of intracellular pantothenate (Figure 2a and Figure 3a). While pyruvate depletion and reduced lactate secretion are likely direct consequences of PDH activation, an increase in the total CoA pool and a concomitant depletion of pantothenate hints at an unexpected activation of *de novo* CoA biosynthesis.

**Figure 3.**
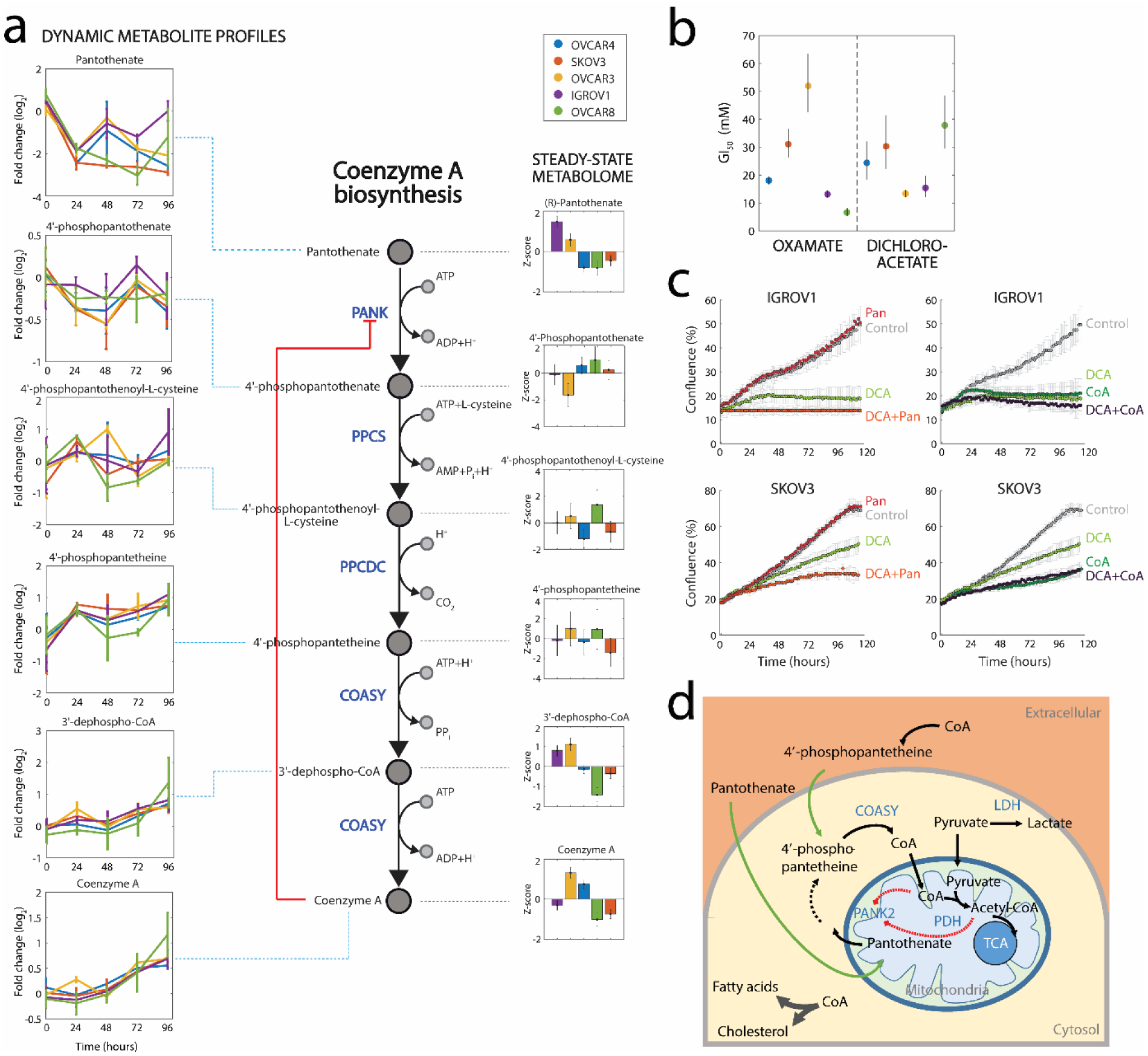
Influence of CoA metabolism on dichloroacetate action. **(a)** Schematic representation of the CoA biosynthetic pathway. Pantothenate kinases (PANK) catalyze the rate-limiting step in CoA biosynthesis^70^. For each pathway intermediate, relative steady-state abundances in the untreated condition (right-hand side) and dynamic changes upon dichloroacetate treatment (left-hand side) are shown. **(b)** Determined GI_50_ concentrations for oxamate and dichloroacetate across cell lines (Supplementary Figure S10). **(c)** IGROV1 and SKOV3 cells were grown in RPMI1640 medium before addition of perturbing agents and continuous confluence monitoring for approx. 5 days. Six conditions were tested: normal RPMI1640 medium (Control), addition of 2.1 μM pantothenate, with and without 11 mM (IGROV1) or 25 mM (SKOV3) of dichloroacetate (Pan/Pan+DCA), addition of 100 μM CoA with and without 13 mM (IGROV1) or 31 mM (SKOV3) of dichloroacetate (CoA/CoA+DCA). **(d)** Schematic representation of CoA metabolism. CoA plays a central role in energy and fatty acid metabolism, acting as an acyl group carrier to form acetyl-CoA and other important compounds, such as fatty acids, cholesterol and acetylcholine. PANK2, the first and rate-limiting metabolic enzyme in the CoA biosynthetic pathway, is allosterically regulated^41,43^ and localizes in the mitochondrial inter-membrane space^40–43,71^. CoA is produced in the cytosol and subsequently actively transported into the mitochondrial matrix. Alternatively, cells are able to scavenge CoA from the extracellular environment thanks to the action of extracellular ectonucleotide pyrophosphatases contained in the serum^72^. These enzymes cleave the CoA molecule to form 4’-phosphopantetheine, which can enter the cells one enzymatic step above CoA formation by COASY.

Pantothenate is the primary precursor required for CoA biosynthesis. CoA in turn regulates its own biosynthesis via allosteric inhibition of the first enzymatic step in the pathway, catalyzed by mitochondrial pantothenate kinase 2 (PANK2)^40,41^ (Figure 3a,d). While human PANK2 locates in the inner mitochondrial membrane^42^, the remaining CoA biosynthetic steps take place in the cytoplasm. Notably, CoA pools in the different cellular compartments are tightly regulated, such that typical CoA concentrations are 1-2 orders of magnitude higher in mitochondria (∼2-5 mM) than in the mitochondrial intermembrane space and the cytosol (∼0.02-0.14 mM)^43,44^. We hypothesize that a hyper-activation of PDH in the mitochondrial matrix could entail a depletion of CoA in the immediate surroundings of PANK2, hence lifting allosteric inhibition of *de novo* CoA biosynthesis. In such a scenario, our observations are consistent with an attempt of cells to re-equilibrate CoA levels across compartments upon dichloroacetate treatment by activating the CoA biosynthetic pathway (Figure 3a). According to this model, the resulting increased pantothenate phosphorylation would explain the observed depletion of intracellular pantothenate, and the parallel accumulation of total CoA in the cells^45^ (Figure 3a). As result, the imbalance between cytosolic production and dicholoroacetate-induced over-consumption of CoA in mitochondria could be at the basis of dicholoroacetate toxicity. Moreover, we observed the largest amount of intracellular pantothenate at steady state (Figure 3a) in the two cell lines with the highest sensitivity (i.e. lowest GI_50_) to dichloroacetate, OVCAR3 and IGROV1 (Figure 3b and Supplementary Figure S10), supporting a functional association of CoA metabolism with the MoA of dichloroacetate.

To test our hypothesis, we selected IGROV1 and SKOV3 cells, which exhibit different steady-state levels of pantothenate and a distinctly different sensitivity to dichloroacetate (Figure 3b). We monitored the growth of IGROV1 and SKOV3 cells upon dichloroacetate treatment, with and without supplementing the medium with 2.1 μM of pantothenate or 100 μM of CoA (Figure 3c). The results were twofold: (i) increasing extracellular pantothenate concentration strongly aggravated the toxicity of dichloroacetate in both cell lines (Figure 3c), while being slightly beneficial for cells in normal RPMI1640 medium (containing 0.25 μM pantothenate), (ii) surprisingly, even in the absence of dichloroacetate, supplementing CoA to the medium had a strong toxic effect on cells. Cells supplemented with 100 or 500 μM CoA (Figure 3c and Supplementary Figure S11) exhibited a first phase of normal growth, followed by rapid growth arrest (Figure 3c). Interestingly, supplementation of CoA completely masked dichloroacetate toxicity when co-administered (Figure 3c and Supplementary Figure S11). The synergistic effect of pantothenate with dichloroacetate, and the antagonistic interaction of dichloroacetate with CoA reinforce our premise of a functional interplay between CoA metabolism and cell growth inhibition caused by dichloroacetate (Figure 3d). Because CoA biosynthesis is regulated (i.e. repressed) immediately downstream of pantothenate (Figure 3a), CoA levels can be controlled in spite of high pantothenate concentrations. Hence, supplementing pantothenate to the medium has no toxic effect to cells (red curve in Figure 3c). However, when cells are challenged with dichloroacetate, CoA biosynthesis is activated and higher levels of pantothenate can lead to higher CoA biosynthetic flux, which in turn aggravates dichloroacetate toxicity (orange curve in Figure 3c). Conversely, by directly providing CoA extracellularly, we bypassed the possibility for cells to control CoA homeostasis, inducing high toxicity. Taken together, our experimental findings suggest the dichloroacetate toxicity to be directly linked to CoA overproduction.

We next asked whether CoA toxicity was restricted to ovarian cancer cell lines. To this end, we tested the effect of CoA on eight additional cancer cell lines from different tissue types. Despite distinct differences in sensitivity, all cell lines exhibited growth reduction upon supplementation of culture media with 100 μM CoA (Figure 4). To our knowledge, this is the first time that a toxic effect of CoA in mammalian cells has been shown and was linked to the mode of action of dichloroacetate. While outside the scope of this work, CoA toxicity might hence deserve more attention in future studies. Consistent with our *in vitro* results, *in vivo* inhibition of PANK in mice by hopantenate, a competitive inhibitor of pantothenate kinase, resulted in 167-fold higher expression of PDK in a previous study^46^. This observation suggests dichotomous compensatory mechanisms to regulate CoA homeostasis: (A) inhibition of CoA biosynthesis activates PDK, which in turn represses PDH^46^, while (B) inhibition of PDK by dichloroacetate has the opposite effect, and promotes CoA biosynthesis. The resulting over-induction of cytosolic *de novo* CoA biosynthesis and accumulation of CoA can in turn aggravate dichloroacetate toxicity. To test whether the observed adaptive response was an indirect effect associated with a general stress response upon growth inhibition and/or reduced lactate secretion, we tested the effect of oxamate, a small molecule that inhibits the conversion of pyruvate into lactate.

**Figure 4.**
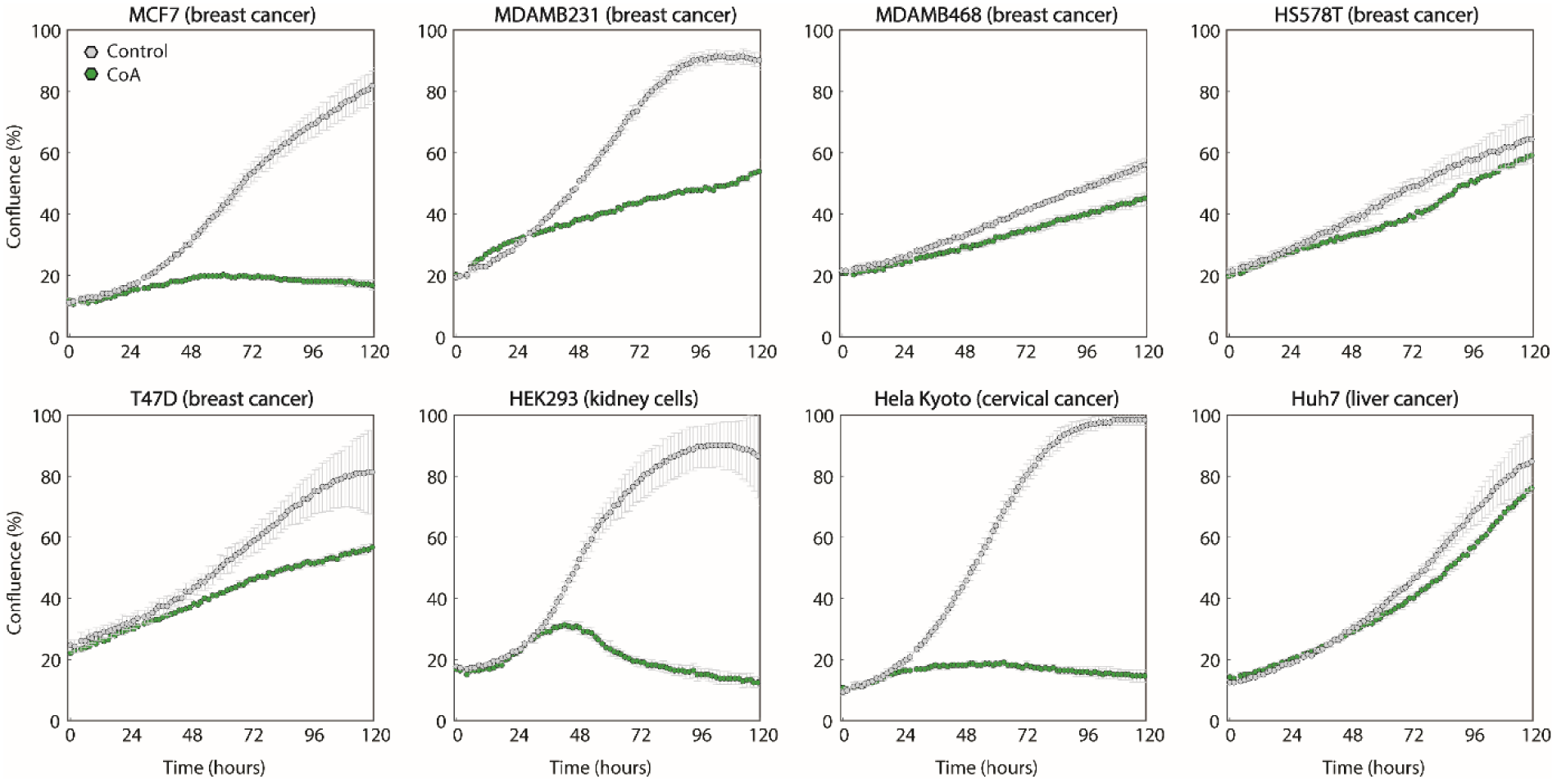
CoA toxicity across multiple cell lines. Cell lines from breast, kidney, cervical and liver tissues were grown with (green) and without (grey) the addition of 100 μM CoA. Cell confluence is reported as mean ± SD across three replicates.

### Oxamate and dichloroacetate elicit different metabolic responses

Oxamate is a competitive inhibitor of lactate dehydrogenase (LDHA) with respect to pyruvate^47^ (Figure 2b). A growing body of evidence indicates that oxamate induces apoptosis exclusively in cancer cells^47,48^. According to current theory, the inhibition of lactate production, together with typically high glycolytic rates in cancer cells, causes an over-production of toxic superoxide by the mitochondrial electron transport chain. Since both drugs decrease lactate secretion rates (Supplementary Figure S8), oxamate treatment could lead to similar metabolic adaptive mechanisms as dichloroacetate.

In our dynamic metabolome profiling data, we observed a significant accumulation of intermediates in TCA cycle upon oxamate treatment, and a concomitant reduction of intracellular ATP levels, in accordance with previous findings^49^. Overall, we found that the significant metabolic changes common to all cell lines locate in central metabolic pathways like oxidative phosphorylation and nucleotide metabolism (Figure 2c). In particular, we observed a consistent and large accumulation of sorbitol and sedoheptulose 7-phosphate (Figure 2b). Overall, changes induced by oxamate were largely different from those induced by dichloroacetate (Figure 3a-c), suggesting for radically different metabolic adaptive strategies. Unlike PDH activation by dichloroacetate, inhibition of lactate dehydrogenase seems to redirect intermediates in upper glycolysis to other pathways, such as NADPH-dependent reduction of glucose to form sorbitol, or the pentose phosphate pathway (as indicated by accumulation of sedoheptulose 7-phosphate, Figure 2b). Both metabolic responses are known to counteract oxidative stress^50,51^. Interestingly, we also observed a marked reduction in the level of *N*-Acetylaspartic acid (Figure 2b), a potent oxidative stress agent^52^ associated with poor prognosis in ovarian cancer^53^. The marked differences between metabolic adaptive responses to oxamate and to dichloroacetate reinforce our previous observation of a selective functional link between CoA metabolism and the mode of action of dichloroacetate.

## Conclusions/Discussions

In this study, we present a novel experimental and computational workflow that enables a systematic and high-throughput investigation of dynamic changes in the intracellular metabolism of adherent mammalian cells upon environmental perturbation. Our methodology provides a novel way to perform high-throughput dynamic metabolic screens in adherent cell lines, and addresses several challenges in computational data normalization and interpretation. Limitations concern both the extraction of the most informative features from non-targeted profiling data, and the general ability to systematically infer testable molecular hypotheses from non-targeted metabolomics screens. Our approach offers a higher throughput than previous state-of-the-art metabolomics methodologies for adherent mammalian cells^27–28,54,55^ by using a miniaturized parallel 96-well cultivation system, a simple metabolite extraction procedure and automated time-lapse microscopy. We additionally exploit the unique advantages of flow-injection high-resolution MS-based metabolomics that provides exceptional throughput and repeatability, and has become an invaluable tool^56–59^ for exploratory studies and the profiling of large sample collections. Altogether, our methodology offers new scientific and clinical opportunities for large-scale *in vitro* exploratory metabolome drug screenings and a complementary tool to more targeted approaches^60^.

A major challenge common to many high-content screenings is the computational analysis of large datasets for the generation of testable predictions. Here, we implemented a systematic data processing and analysis pipeline that allows comprehensively interpreting dynamic metabolic profiles (Supplementary Figure S2). While changes in metabolite abundance do not necessarily correspond to changes in conversion rates (i.e. fluxes), altered metabolite pools can be reflective of functional changes in the cell^61^. By investigating the dynamic responses to the perturbing agents dichloroacetate and oxamate, we proposed a previously undescribed role of pantothenate and CoA in mediating the toxicity of dichloroacetate. Therapeutically, high dosages of dichloroacetate are needed in order to effectively suppress tumour growth^37^, limiting further development and usage of this compound in clinics. Nevertheless, our results suggest that compounds affecting CoA production are likely to exhibit strong epistatic interactions with dichloroacetate. Combining dichloroacetate with activators of PANK enzymes^62^ could hence potentially increase the therapeutic efficacy and allow reducing the dichloroacetate dosage, offering an attractive approach to reduce dichloroacetate side effects. In light of the promising initial evidence that we provided here, this possibility warrants more attention in future studies.

We have shown here that our experimental and computational framework for high-throughput drug metabolome profiling can provide key insights into the cellular response to bioactive compounds. As such, this technique can become a powerful complementary tool to aid lead selection at early stages of drug discovery, and to predict compound modes of action, similar to approaches exploiting large compendia of cellular gene expression profiles ^21^. For instance, comparative analysis can reveal uncharacterized compounds featuring metabolic responses similar to drugs with known molecular targets^2^. Our proof-of-principle example illustrates how *in vitro* high-content metabolic drug profiling can provide a first coarse-grained characterization of a compound mode of action. It is worth noting that the here presented methodology is widely applicable to more clinically relevant models, like primary cells, and can be a powerful tool in guiding the rational design of *in vivo* low-throughput follow-up studies. Despite the difficulty in translating the relevance of *in vitro* phenotypes into *in vivo* outcomes^63^, we envisage that this approach can be applied to the profiling of large sets of bioactive compounds^31^ in a large cohort of cell lines^64,65^. In such a setting, this methodology can potentially deliver invaluable insights to (i) highlight mechanistic biomarkers to be tested *in vivo*, (ii) resolve the functionality of genetic variations, and (iii) understand the interplay between the drug mode of action and intrinsic cell-to-cell tolerance variability.

## Availability of data and material

All data generated or analysed during this study are included in this published article as supplementary materials.

## Funding / Acknowledgments

We thank the National Cancer Institute (NCI) for providing the cancer cell lines. This work was supported by a Worldwide Cancer Research (WCR-15-1058) project funding to M.Z., K.O. was funded by the Austrian Science Fund (FWF): FWF P26603 and FWF W1224 Doctoral Program BioToP – Biomolecular Technology of Proteins. We thank Uwe Sauer and Nicola Zamboni for supporting this work and providing laboratory facilities, Dimitris Christodoulou, Maren Diether, Duncan Holbrook-Smith and Victor Chubukov for helpful feedback and discussions.

## Authors’ information / Contributions

M.Z. designed the project. S.D. and K.O. performed the metabolome experiments. S.D. and M.Z. designed and implemented the image analysis framework. K.O. performed follow up experiments. M.Z. K.O. and S.D. analyzed the data. All authors contributed to preparing the manuscript.

## Materials and methods

### Cell cultivation

The ovarian cancer cell lines IGROV1, OVCAR3, OVCAR4, OVCAR8 and SKOV3 were obtained from the National Cancer Institute (NCI, Bethesda, MD, USA) and maintained according to standard protocols at 37°C with 5% CO2 in RPMI1640 (Biological Industries, cat. No. 01-101-1A) supplemented with 2 mM L-glutamine (Gibco, cat.no. 25030024), 2 g/L D-Glucose (Sigma Aldrich, cat.no. G8644), 100 U/mL Penicillin/Streptomycin (Gibco, cat.no. 15140122) and 5% fetal bovine serum (FBS, Sigma Aldrich, cat.no. F6178). After thawing, the cells were expanded in standard cell culture flasks (Nunc T75, Thermo Scientific). After one week, the cells were transferred to fresh medium where FBS was replaced by dialyzed FBS (Sigma Aldrich, cat.no. F0392) in order to facilitate metabolite quantification. Cells were maintained in this medium with dialyzed FBS for the remaining duration of the experiment. Overall, the cell lines were expanded in T75 for a total of three weeks from thawing until perturbation and metabolomics experiments. With the aim of determining the starting cell density for the experiment, a preliminary cultivation in 96-well cell culture plates was done one week prior to the experiment. To this end, for each cell line, 150 μL of 8 different dilutions containing different starting cell numbers were plated in triplicate in a 96-well plate. After 72 hours, all wells were imaged using a Spark^™^ 10M (TECAN) and confluence was determined in each well. The optimal starting cell density was subsequently calculated so as to obtain 80% confluence after 72 hours.

### Perturbation experiments

Sodium oxamate and sodium dichloroacetate were obtained from Sigma Aldrich (cat.no. O2751 and 347795, respectively), and stock solutions of 400 mM oxamate and 250 mM dichloroacetate were prepared in distilled water. To determine the GI_50_ drug concentrations, nine different concentrations of oxamate and dichloroacetate were tested (Supplementary Figure S10). For each cell line, cells were seeded in 135 μL of fresh medium in 96-well plates according to the previously calculated optimal density. After 24 h, oxamate and dichloroacetate, dissolved at different concentrations in 15 μL of medium, were added to the cells in triplicates. Immediately upon drug addition, 24 h, 48 h and 72h after drug exposure, all wells were imaged using a Spark^™^ 10M (TECAN) plate reader, and the cell confluence was determined. For each condition and cell line, the growth rate was obtained by fitting an exponential curve to the cell confluence measurements. The relative growth rate inhibition upon different drug concentrations was then fitted by a sigmoidal curve (Supplementary Figure S10) and the drug concentration causing a 50% reduction in growth rate (GI_50_) was estimated from the fitted curve (Figure 3b).

### Metabolomics experiments

Cell lines were plated in nine 96-well plates according to the optimal density previously calculated, using 135 μL of medium. To minimize the effect of evaporation, the outmost rows and columns of the plate were omitted, and filled with PBS instead. After 24 h, cells were perturbed with 15 μL of medium containing drug concentrations close to the respective GI_50_ for each cell line. When the calculated GI_50_ dose could not be reached due to limited solubility, the highest concentration possible was used (400 mM and 250 mM for oxamate and dichloroacetate, respectively). 15 μL of fresh medium without drug addition were used as a control. The final concentrations used for each drug and each cell line are given Table 1.

### Sample collection and metabolite extraction

The metabolomics sampling procedure was adapted from an experimental workflow for steady-state metabolome profiling described in ^29^, and is here briefly summarized. Samples were collected immediately before, and at 24 h, 48 h, 72 h and 96 h after drug addition. Two replicate 96-well plates were processed at each sampling time point (plate A and plate B, see also Supplementary Figure S1). In plate A, the cell culture medium was aspirated from all wells using a multichannel aspirator, and 150 μL of ammonium carbonate (75 mM, pH 7.4, 37°C) was gently added to each well using a multichannel dispensing pipet. Immediately after aspiration of the washing solution, 100 μL of cold extraction solvent (40% methanol, 40% acetonitrile, 20% water with 25 μM phenyl hydrazine^66^, −20°C) were added to each well using a multichannel pipet. Plates were sealed with aluminium adhesive to prevent evaporation, immediately transferred to −20°C for 1 h, and subsequently stored at −80°C until further processing. In plate B, the cell culture medium was aspirated, and the cells were washed with ammonium carbonate (75 mM, pH 7.4, 37°C). After aspiration of the washing solvent, 150 μL of PBS (Gibco, cat.no. 10010015) were added to each well, and cell confluence was immediately measured in all wells using a Spark^™^ 10M (TECAN) plate reader, adopting bright-field microscopy. The cell confluence from plate B was later used as a measure of extracted cell number for normalization. Before injection in the mass spectrometer, the 96-well plates were briefly thawed on ice, and the bottom of all wells was scratched using a multichannel pipet with wide-bore tips in order to disrupt and detach all cells from the well bottom. The plates were centrifuged (4°C, 4000 rcf), and the supernatant was transferred to fresh 96-well plates for FIA-TOFMS measurements.

### FIA-TOFMS analysis

Flow-injection time-of-flight mass spectrometry (FIA-TOFMS) analysis was performed as described in ^25^ on an Agilent 6550 iFunnel Q-TOF LC/MS System (Agilent Technologies, Santa Clara, CA, USA) equipped with an electrospray ion source operated in negative ionization mode. In this setup, the samples are injected into a constant flow of an isopropanol/water mixture (60:40, v/v) buffered with 5 mM ammonium carbonate at pH 9 using a Gerstel MPS2 autosampler (5 μL injection volume). Two compounds were added to the solvent for on-line mass axis correction: 3-Amino-1-propanesulfonic acid, (HOT, 138.0230374 m/z, Sigma Aldrich, cat. no. A76109) and hexakis(1H,1H,3H-tetrafluoropropoxy)phosphazine (940.0003763 m/z, HP-0921, Agilent Technologies, Santa Clara, CA, USA). The ion source parameters were set as follows: 325°C source temperature, 5 L/min drying gas, 30 psig nebulizer pressure, 175 V fragmentor voltage, 65 V skimmer voltage, 750 V octopole voltage. The TOF detector was operated in 4 GHz high-resolution mode with a spectral acquisition rate of 1.4 spectra per second. Mass spectra were recorded in the mass range 50-1000 m/z. Alignment of MS profiles and picking of centroid ion masses were performed using an in-house data processing environment in Matlab R2015b (The Mathworks, Natick)^25^.

### Ion annotation

The ion annotation process is based on a list of known metabolites, compiled from the Human Metabolome Database (HMDB^34^) and the Recon2 genome-scale reconstruction of human metabolism^35^. In order to allow annotation of α-keto acid derivatives formed in presence of phenyl hydrazine^66^ in the extraction solvent, we added the sum formulae for the phenylhydrazones (+C_6_H_8_N_2_ −H_2_O) of a total of 30 α-keto acid compounds (selected via KEGG SimComp search http://www.genome.jp/tools/simcomp) to the metabolite list for annotation. The monoisotopic mass is calculated for each of the listed metabolites based on its sum formula. A list of expected ion masses corresponding to the listed metabolites is subsequently generated, considering only ionization by deprotonation (-H+) in negative mode electrospray ionization. Subsequently, these theoretical ion masses are searched against the detected ion mass-to-charge ratios (m/z) within a tolerance of 0.003 amu. The final list of annotated ions is compiled considering the best metabolite match (i.e. smallest difference to the expected mass) for each ion.

### Data processing and computational analysis for steady state metabolome data

All steps of data processing and further analysis were performed in Matlab 2015b (The Mathworks, Natick). For steady-state metabolome profiles, the bioinformatics pipeline is described in ^29^ and is here summarized. Multiple regression analysis to estimate the relative metabolite concentrations at steady state was performed using the Matlab *fitlm* function. This function infers model parameters α (cell line-specific) and β by minimizing the Euclidian distance between measured metabolite intensities and model predicted ones. It is worth noting that the β represents the MS background signal, or in other words the ion intensity when no cells are extracted. Hence, this particular parameter is independent from cell types. Because of the difficulties in reliably estimating the extracted cell number from bright-field microscopy images above a confluence of 80% (Supplementary Figure S3), and the observed deviation from metabolic steady-state (Supplementary Figure S4), we excluded all metabolome measurements taken above this cell density threshold. For each metabolite, we solve the following linear model:

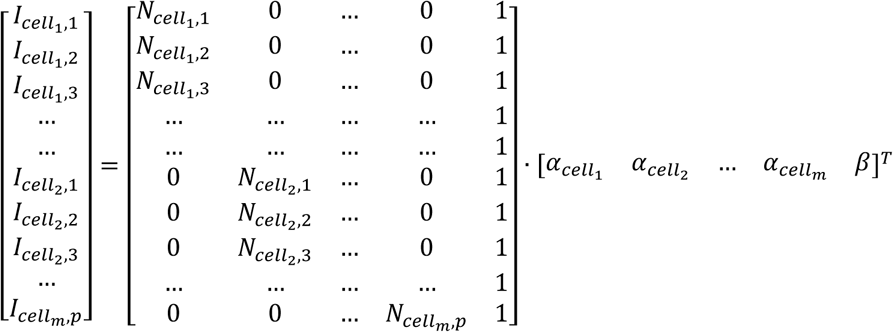

Where I_ce_ll1,1 is the measured metabolite intensity in cell-line_1_/sample_1_, N_cell1,1_ is the corresponding number of cells extracted in cell-line_1_/sample_1_. Cell line specific αs and β are the unknown parameters to be fitted. We selected the metabolites exhibiting a significant alteration in at least one cell line using a one-way ANOVA test, including a step correcting for multiple hypothesis testing^67^–^69^ (Supplementary Figure S12).

### Data processing and computational analysis for dynamic drug-induced metabolome changes

The full matrix of dynamic metabolic profiles after dichloroacetate and oxamate treatments is provided in Supplementary Data Table S2. In order to deduce a specific metabolic fingerprint induced by an external perturbing agent, we (i) selected the most significant metabolic changes conserved across the cell lines, (ii) performed pathway enrichment analysis on the resulting list of metabolites, and (iii) analyzed response variability across all cell lines.

i. Separately for each perturbation (i.e. oxamate and dichloroacetate), we extracted the most significant and prominent metabolic changes that are conserved across the different cell lines. To this end, for each individual metabolite time course we calculated the median of maximum absolute fold changes and the product of lowest p-values across cell lines. As a result, each metabolite is associated with a unique median fold-change and p-value, summarizing the effect of the perturbation on all cell lines.
ii. Metabolites with an absolute log_2_ fold-change **≥** 1 and a combined p-value **≤** 1e-10 are tested against KEGG metabolic pathways. Pathways with an overrepresented number of altered metabolites are selected based on a hypergeometric statistical test and p-value correction for multiple tests^67,68^. (iii) Metabolites that exhibit cell line-specific responses to a given perturbation are selected on the basis of the response variability exhibited across the different cell lines. The standard deviation for each metabolite was calculated from the aforementioned maximum fold changes in each cell-line time course, and metabolites with a standard deviation ≥ 1.5 are retained and subjected to pathway enrichment analysis (Supplementary Figure S12).

### Cell growth and segmentation

All procedures for cell growth monitoring and image analysis were adopted from ^29^, and are here briefly summarized. The monitoring of cell growth during the entire course of the experiment was performed using A TECAN Spark 10M plate reader was used to monitor live adherent cell cultures directly in the 96-well culture plate. The choice of image acquisition frequency depends on how fast are the expected growth dynamic changes. Here, we selected a time frequency of 1.5 hours as a reasonable tradeoff between the fastest doubling time among our cell lines (approx. 20 hours) and the time it takes to acquire the images for a full plate on the TECAN plate reader (approx. 30 min). It is worth noting that our procedure can be adapted to other commercially available plate readers. Full detail on bright-field image processing and the extraction of cell confluence and average adherent cell size is described in ^29^ (MATLAB code available for download).

